# Replacement of a single residue in an antibody completely abolishes cognate antigen binding, as predicted by theoretical methods

**DOI:** 10.1101/2025.06.26.661714

**Authors:** Marvin Scherlo, Adrian Höveler, Marvin Mann, Grischa Gerwert, Jörn Güldenhaupt, Klaus Gerwert, Till Rudack, Carsten Kötting

**Affiliations:** Center for Protein Diagnostics (PRODI), Biospectroscopy, Ruhr University Bochum, Bochum, Germany; Department of Biophysics, Ruhr University Bochum, Bochum, Germany; Biomolecular Simulations and Theoretical Biophysics Group, Faculty of Biology and Biotechnology, Ruhr University Bochum, Bochum, Germany; Structural Bioinformatics Group, Regensburg Center for Biochemistry, University Regensburg, Regensburg, Germany; Structural Bioinformatics Group, Regensburg Center for Ultrafast Nanoscopy, University Regensburg, Regensburg, Germany

**Author notes:** these authors contributed equally (Marvin Scherlo, Adrian Höveler).

## Abstract

Structural insights into the interaction between antibodies and antigens at the atomic level are pivotal for understanding the molecular mechanisms of antigen binding. Despite the availability of structural models generated by recent artificial intelligence advancements, computational predictions require experimental validation to confirm their accuracy. Here, we demonstrate a novel approach that combines computational protein modeling with spectroscopic experiments to validate antibody-antigen interactions. As a case example we use solanezumab, a monoclonal antibody that targets amyloid-beta (Aβ), whose misfolding is the main factor responsible for Alzheimer’s disease. For this antibody we predicted a single mutation, G95A^HC^, within the paratope of the heavy chain to disrupt antigen binding. This mutation, referred to as a “dead mutant”, was experimentally validated using an immuno-infrared biosensor (iRS). Our results confirmed that the mutation abolished antigen binding without affecting the native structure of the antibody. The use of dead mutants enables precise differentiation between specific and nonspecific binding, which is particularly important in medical diagnostics. We applied this approach to analyze the binding of solanezumab to synthetically produced Aβ variants and Aβ catched by the iRS functionalized surface from cerebrospinal fluid, showcasing its utility in Alzheimer’s disease diagnostics. These findings highlight the value of computational modeling and experimental validation in understanding antigen-antibody interactions, with significant implications for diagnostic and therapeutic applications.

## Introduction

For a comprehensive understanding of the interaction between antibodies and their corresponding antigens, structural information at the atomic level is crucial. However, experimentally resolved structural models of antibodies with bound antigens are scarce. Owing to recent artificial intelligence-driven advances in structure prediction, many such models are becoming easily accessible ^1^. However, both predicted and experimentally resolved models create a demand for analysis tools to derive key interaction patterns from the models to discover the molecular mechanism underlying antigen binding. The computationally identified interaction mechanism is still hypothetical and requires experimental validation ^2^. A versatile tool for this purpose is site-directed mutagenesis. A small alteration within a theoretically predicted interaction site should drastically influence the dissociation constant between the antibody and the antigen. Ideally, this alteration will result in a substantial reduction or elimination of the interaction, creating what is referred to as a “dead mutant”.

Thus, experimental validation of a dead mutant provides direct knowledge about the antigen-antibody interaction. This can in turn be used to further refine theoretical models for a detailed knowledge of the binding characteristics.

Dead mutants serve as powerful tools in various applications, particularly in diagnostics, where they function as ideal negative controls. For example, in a sensor based on the binding of the target to an antibody, the same antibody with an impaired binding site would prevent specific binding. This allows the study of unintended binding, e.g. at glycosylation sites. Alternatively, a sensor surface without an antibody would not show unintended binding to the antibody and could have very different properties with respect to unspecific binding on the surface. Thus, during assay development, the use of dead mutants enables the precise differentiation between specific binding at the antigen binding site, unintended binding at other sites of the antibody and nonspecific binding, enhancing assay specificity and reliability.

Immuno-infrared biosensors (iRS) are based on attenuated total reflectance Fourier transform infrared (ATR-FTIR) spectroscopy ^3^. They operate label-free and thereby offer significant advantages over many other types of biosensors ^4^. In addition to the usual binding information, these sensors provide information on the secondary structure distribution of the antigen. This feature is particularly advantageous for studying proteinopathies, making it a versatile diagnostic tool. In the case of Alzheimer’s disease, all Aβ from a CSF or blood sample is immobilized. The more misfolded Aβ with high β-sheet content present, the lower the position of the amide I absorption measured in the infrared spectrum. This powerful method is applicable to diagnose Alzheimer’s disease many years before clinical symptoms manifest ^5–7^. However, a limitation of label-free infrared spectroscopy is its inability to distinguish between specific interactions at the binding site and potential nonspecific binding events, as all immobilized proteins contribute equally to an integrated signal. Here, the use of a dead mutant becomes crucial, serving as an ideal control for unequivocally identifying specific interactions at the antigen binding site, thereby enhancing the diagnostic utility of the assay.

Solanezumab was one of the first therapeutic antibodies for Alzheimer’s diseases to enter phase 3 clinical trials ^8^. While solanezumab eventually failed, similar approaches with other antibodies were successful. Today, Lecanemab ^9^ and Donanemab ^10^ are the best therapeutic antibodies for Alzheimer’s disease and are already available as approved drugs. Many other candidates are currently in clinical trials ^11^. A problem for therapy is that diagnosis is usually only made after clinical symptoms, late in the overall progression of the disease. Early intervention is expected to be key to the successful treatment of Alzheimer’s disease ^12^. iRS have shown its ability to identify Aβ misfolding many years before clinical symptoms occur. Solanezumab is a candidate as a diagnostic antibody for this method.

In this study, we demonstrate the powerful combination of computational protein modeling and spectroscopic experiments at the case example of the monoclonal antibody solanezumab, which targets the Aβ peptide ^8^. In this case, a structural model derived from X-ray crystallography is available^13^. We analyzed the antigen binding site by transferring computational interaction tools initially developed for dynamic interaction analysis within molecular dynamics simulations ^14^ to the analysis of static X-ray structures. Based on the identified antigen-antibody interaction pattern, we propose a single mutation expected to disrupt the interaction. To verify the impact of this mutation, we employed the iRS platform ^15,15–17^. Our results confirmed that the mutation effectively prevents antigen binding, while the native conformation of the antibody remains intact. The interdisciplinary strategy we present here demonstrates the potential of theoretically derived insights to improve biotechnological experimental setups used for disease diagnosis and therapy, thereby narrowing the gap between basic science and clinical applications.

## Results and Discussion

### Theoretical derivation of a dead mutant candidate

The basis for the mutational analysis was the crystal structure of the Aβ mid-domain captured by the solanezumab antigen binding fragment (Fab) (PDB-ID: 4XXD), with a resolution of 2.41 Å ^13^. Our contact analysis revealed that the central binding motif of the antigen, consisting of F19 and F20 of Aβ, is deeply buried in the binding pocket between the antibody heavy and light chains (Fig. 1 A-C). Hydrophobic interactions of the neighboring L17 also strongly contributes to binding, while backbone hydrogen bonds of F19 and A21 to S91^LC^ (LC = Light Chain) as well as the backbone of L17 to D96^HC^ (HC = Heavy Chain) complete their integration into the binding pocket. Several polar residues flank this antigen core region, most notably K16 of Aβ, forming a salt bridge with the side chain of D96^HC^, and D23 of Aβ h-bonded with the side chain and backbone of S33^HC^ (Fig. 1B).

**Figure 1.**
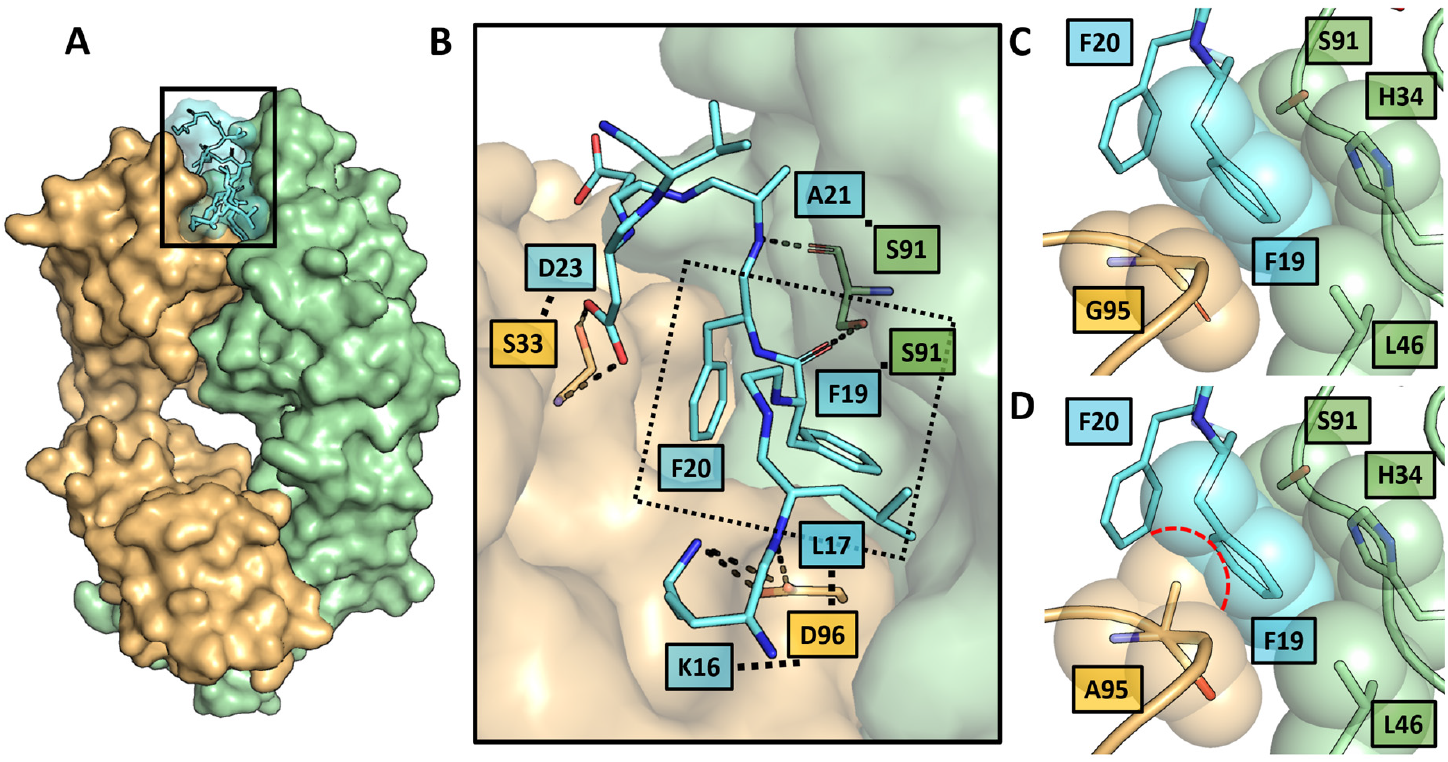
Aβ-peptide binding site of solanezumab wild-type and proposed dead mutant. **A** Structural representation of the solanezumab Fab domain (heavy chain orange, light chain green) with bound Aβ-peptide residues 16-26 (cyan) based on the X-ray structure with PDB-ID 4XXD ^13^. **B** Zoom on the binding site with highlighted hydrogen bond network (dashed lines) of the antibody-antigen interface. **C** highlights the proposed key interaction pattern of F19 (Aβ) buried in a hydrophobic pocket formed by the heavy chain residue G95^HC^ and the light chain residues H34^LC^, L46^LC^ and S91^LC^. **D** Structural prediction of the proposed dead mutant G95A^HC^ illustrating the overlap of the introduced Cβ atom of A95^HC^ with F19 of a bound Aβ-peptide (red dashed lines). Thus, we anticipated that this variant prevents Aβ-peptide binding due to steric hindrance of F19. As we identified this contact as the central interaction motive, we predicted that the loss of this interaction will lead to destabilization of the entire antigen binding motive and thus to a significantly reduced or completely abolished binding affinity.

The asymmetric unit cell comprises two copies of the complex, which superimpose with minimal differences (Cα RMSD of 0.21 Å). The primary distinction is observed for residues G25 and S26 of Aβ: these residues are resolved in chain C, where they engage in crystal packing contacts with symmetry-related molecules, but are disordered and not resolved in chain F in the absence of such stabilizing interactions. As G25 and S26 do not contribute to the antigen-antibody interface, they are omitted from Fig. 1 for clarity. This highlights the importance of considering crystal packing, an often overlooked aspect of theoretical structural studies.

Taking a detailed look on the central double phenylalanine motif, the ring face of F19 is positioned directly above the G95^HC^ Cα atom while the backside is formed by H34^LC^ and S91^LC^, creating a tightly packed binding pocket (Fig. 1C). The ring edges of F19 are further stabilized by close contacts with neighboring residues, including F20 and L46^LC^ among others.

To generate a dead mutant, we aimed to disrupt this central binding motif by substituting G95^HC^ in the CDR H3 region with alanine (Fig. 1C+D). Given the compact nature of the F19 binding pocket, even the minimally invasive introduction of the A95^HC^ methyl group was predicted to prevent F19 from properly inserting into the pocket and to destabilize the entire binding motif. We anticipated that this effect would apply to all tested antigens, as they share an identical central binding motif (Fig. S1).

### Experimental confirmation of the dead mutant

To evaluate the theoretical predictions, we recombinantly produced both the wild-type solanezumab and its dead mutant as Fab conjugated to the “Immunity protein 7” (Im7) using an *E. coli* expression system, as detailed in the Methods section ^18^. The IM7 tag allows for high affinity, reversible binding to DNase domain of colicin E7 (E7) ^19^. For this purpose, the surface of the ATR-crystal was coated with E7 ^20^. The spectra of the surface-bound solanezumab variants are shown in Fig. 2. The similarity of the spectra of the wild-type and the mutant proteins indicates that the mutation does not interfere with the overall structure of the antibody fragment. Structural changes would lead to a change in the shape of the amide I band ^21^.

**Figure 2:**
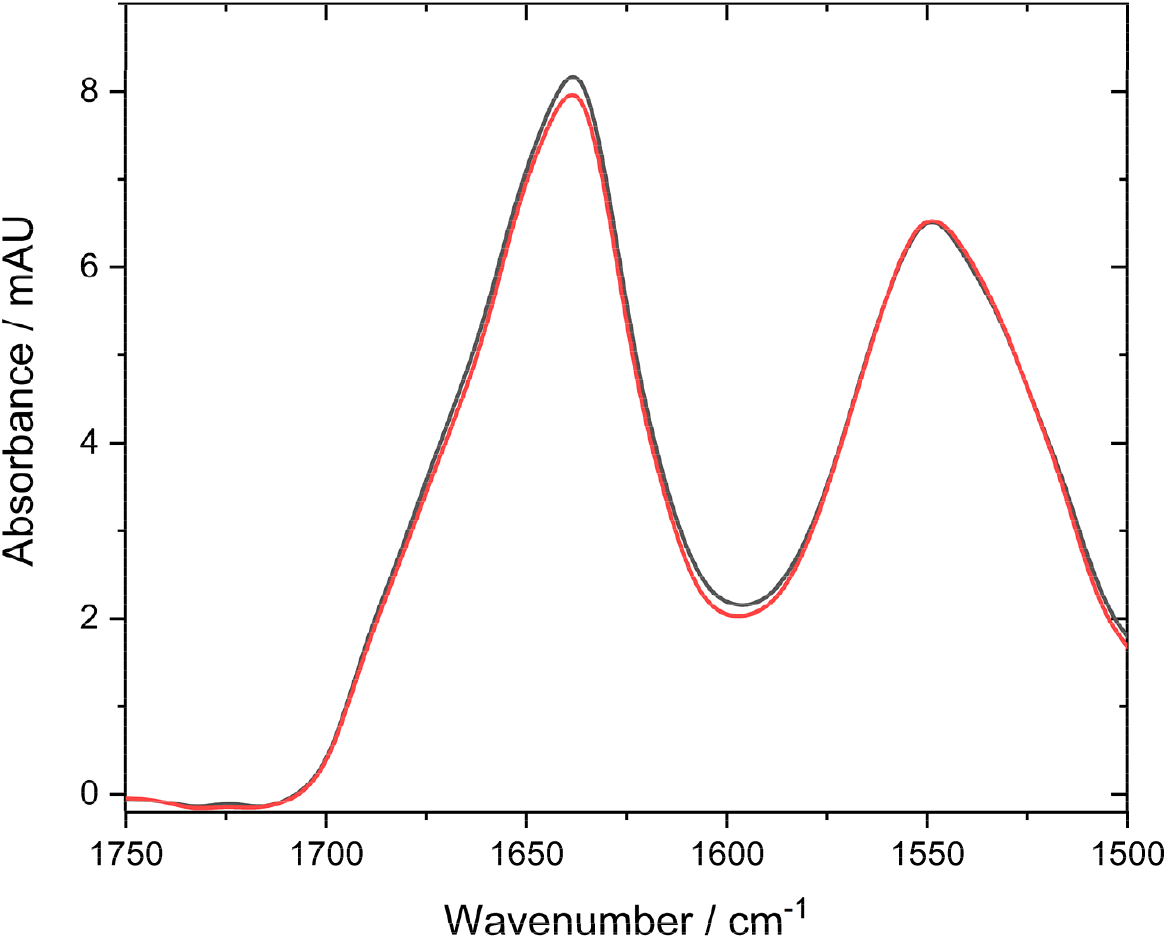
ATR-FTIR spectra of immobilized Solanezumab-Im7. Solanezumab wild-type is shown in black and Solanezumab G95A^HC^ in red. The high similarity of the two spectra indicates the non-invasive nature of the introduced mutation for the structure of the antibody.

Furthermore, we generated several antigen variants (Fig. 3A). The Aβ_9-33_ A21C A30C double mutant is recognized as a monomeric model compound ^22^. The introduction of a disulfide bond reduces the conformational space and impairs fibrilization.

**Figure 3.**
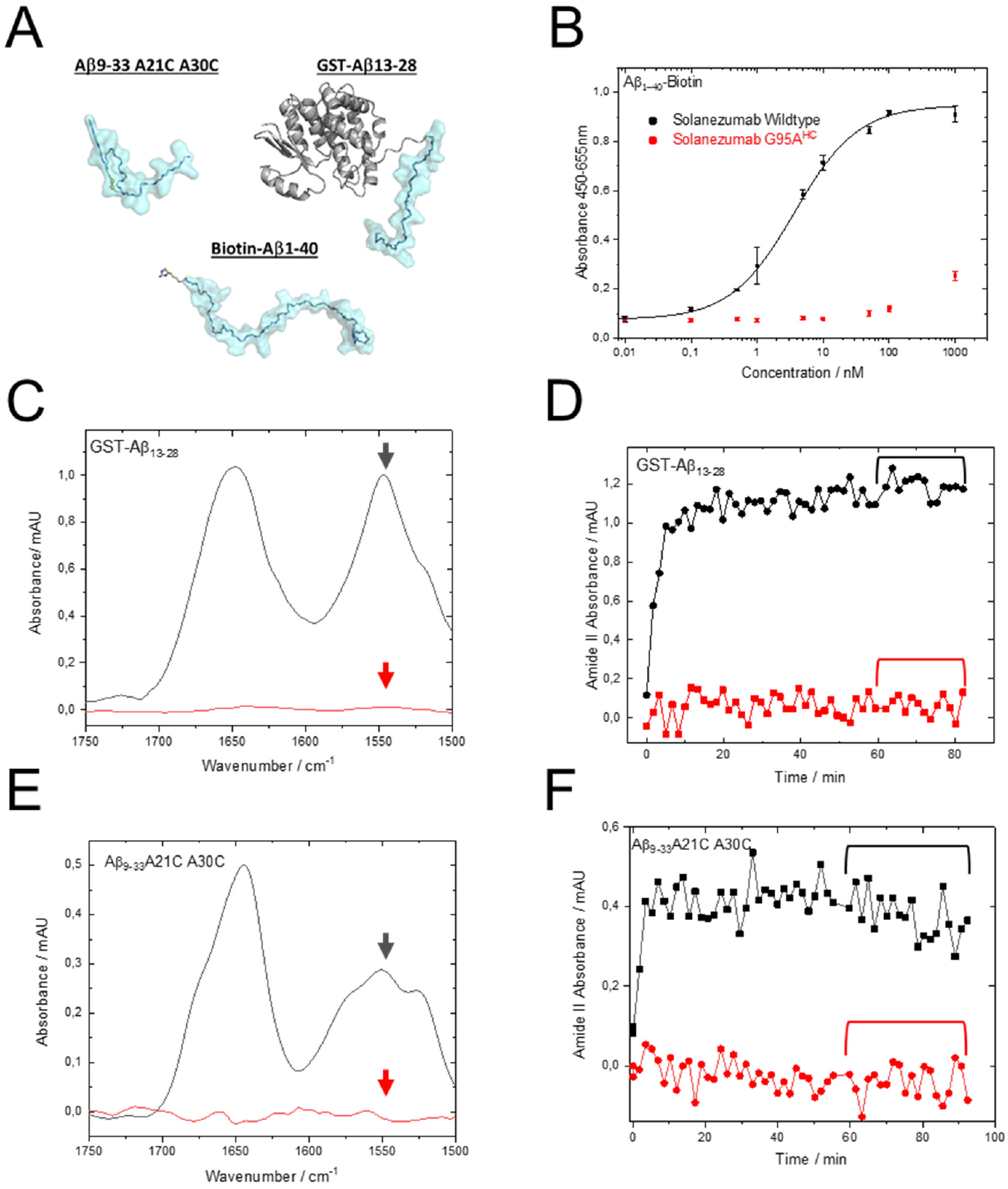
Experimental validation of proposed solanezumab dead mutant. **A** Accurate scale representation of structural models of the three antigen constructs (Aβ cyan, rest gray) investigated. **B** ELISA results demonstrating the binding affinity of soluble biotinylated Aβ1-40 to surface attached solanezumab wild-type (black) and the solanezumab G95A^HC^ dead mutant (red), which was predicted by structural analysis and molecular dynamics simulations. Sigmoidal curves were fitted to the data whenever possible, and the EC50 value was subsequently calculated. **C** ATR-FTIR measurements of the artificial GST-Aβ_13-28_ antigen on a surface coated with solanezumab wild-type (black) and its dead mutant (red). The spectrum of the amide I and amide II region (1700-1500 cm^−1^) is displayed. The wild-type protein exhibits binding of the antigen, with a spectral peak at 1650 cm^−1^, indicating high alpha-helical protein content, while the dead mutant shows no binding. Arrows indicate the amide II region, further elucidated over time in Fig. 3D. **D** Binding kinetics of the amide II band of GST-Aß_13-28_ over time. Protein binding in the wild-type is rapid and reaches saturation. After one hour of sample circulation, the bound antigen is washed (indicated by the bracket), with no observable wash-off. In contrast, the dead mutant shows no protein binding over time. **E** ATR-FTIR measurements of the double cysteine mutant Aβ_9-33_ A21C A30C on a surface coated with solanezumab wild-type (black) and its dead mutant G95A^HC^ (red). Similar to GST-Aß_13-28_ (Fig. 3C), only solanezumab wild-type exhibits binding of the sample and a resulting alpha-helical protein spectrum. The amide I maximum is observed at approximately 1646 cm^−1^. **F** Binding kinetics of the amide II band of Aβ_9-33_ A21C A30C over time. The wild-type (black) rapidly binds the sample in the initial minutes of circulation and gradually saturates. During the wash step (indicated by the bracket), a slight wash-off of the sample over time is observed. Conversely, the dead mutant (red) shows no protein binding over time. Using a variety of methods and antigens, we have shown that the solanezumab dead mutant lacks the ability to bind Aβ in all cases.

GST-Aβ conjugates are produced to enhance the infrared signal of the bound antigen. The absorption coefficient of the amide II band is proportional to the number of amide bonds present ^23^. Aβ 40 contains 39 such bonds, whereas GST-Aβ40 contains 273, leading to a seven-fold increase of the IR signal. Additionally, we generated analogous conjugates with Aβ fragments to facilitate characterization of the binding motif. Given that solanezumab is known to bind to the mid-region of Abeta, we used the GST-Aβ_13-28_ conjugate in our experiments.

First, we characterized the antibodies via ELISA ^24^, as described in the Methods section. Briefly, Aβ-biotin was immobilized on the surface of a commercially available streptavidin-coated plate. The wells were then incubated with varying concentrations of solanezumab. The amount of bound solanezumab was detected via an HRP-conjugated anti-human antibody, followed by HRP-mediated oxidation of o-phenylenediamine (OPD). The results are presented in Fig. 3B. While wild-type solanezumab exhibited a half maximal effective concentration (EC50) of 3.5 nM, no specific binding was detected for the mutant variant. These findings highlight the impressive impact of the single mutation on antibody binding.

In the following experiments, we utilized our iRS setup to characterize the binding of antigen. The binding of antigen was monitored by measuring the absorption at 1550 cm^−1^ (indicated by the arrow in Fig. 3C). This corresponds to the amide II band, which is proportional to the number of amide bonds at the surface and thus to the amount of surface-bound protein. The kinetically resolved absorption data (Fig. 3D) clearly demonstrated rapid binding of the GST-Aβ_13-28_ conjugate to wild-type solanezumab, which reached equilibrium within minutes. After 60-minute incubation, the surface was washed with buffer. The signal remained unchanged, indicating a strong interaction between the antigen and the antibody. In contrast, no binding was observed for the mutant antibody. The corresponding IR spectra are shown in Fig. 3C, where we present the mean spectrum recorded during the wash, as illustrated in Fig. 3D. The position of the amide I band maximum provides insights into the secondary structure of the surface-bound protein. In this case, the amide I band is dominated by the GST moiety, which primarily adopts a helical secondary structure (56 % α-helical, 22 % unordered) ^25^, resulting in an absorption maximum at 1650 cm^−1^.

Fig. 3E and 3F present a similar experiment conducted with Aβ_9-33_ A21C A30C as the antigen. Consistent with previous observations, rapid and stable binding was detected with the wild-type solanezumab, whereas no binding was detected with the mutant variant. The IR spectrum of Aβ_9-33_ A21C A30C exhibited a maximum at 1646 cm^−1^, suggesting minimal β-sheet content. These findings align with the results from protein modeling and MD simulations, which indicate highly similar binding modes for Aß40 and the Aβ_9-33_-33 A21C A30C monomer mutant (Fig. 4 and S4+5).

**Figure 4.**
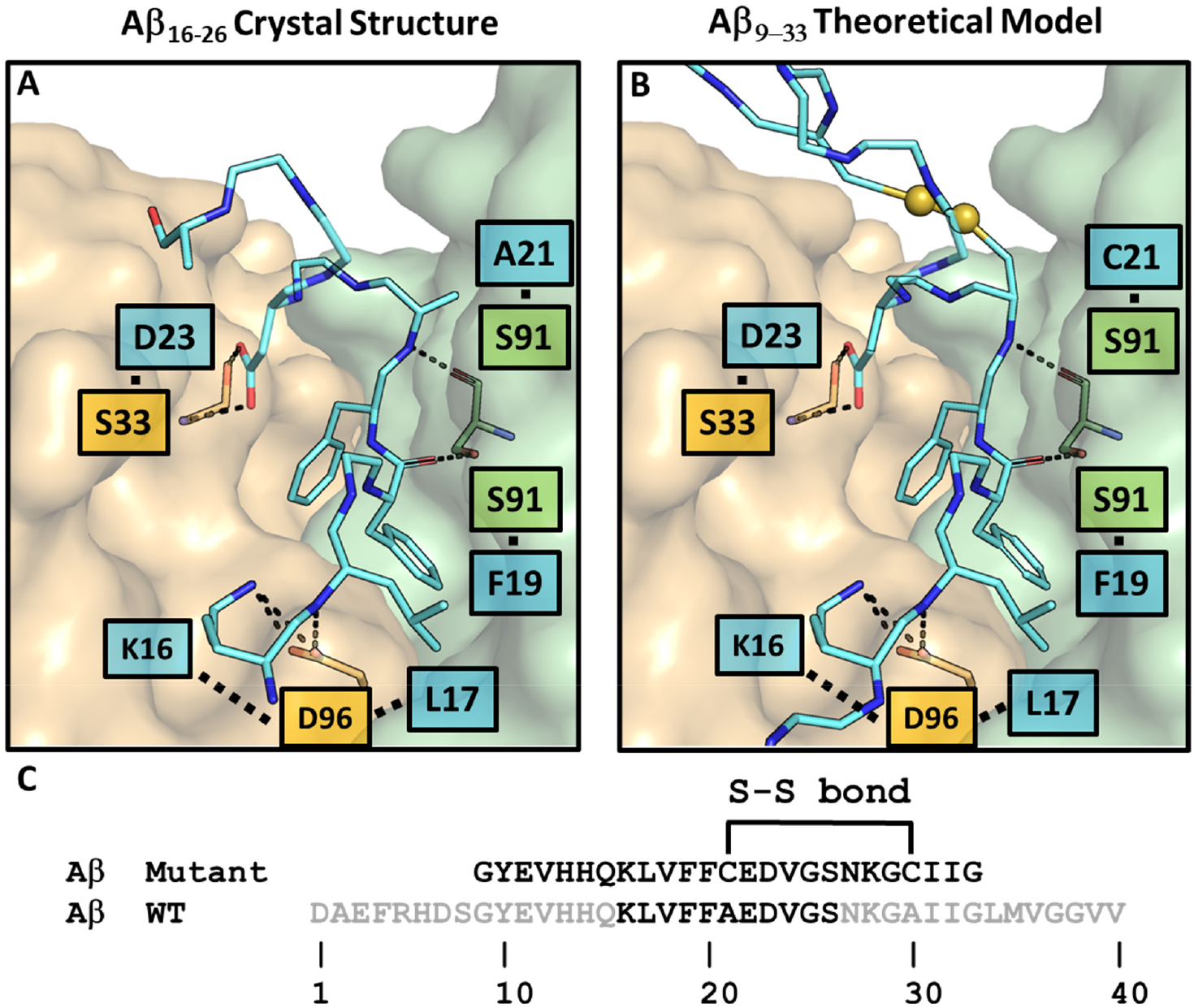
Antigen comparison of Aβ wild-type and Aβ_9-33_ A21C A30C monomer variant. **A** Structural model of the antigen (sticks representation) and antibody (surface representation, heavy chain: orange, light chain: green). The binding interface of Aβ 16-26 wild-type (carbon atoms cyan, nitrogen atoms blue, oxygen atoms red, sulfur atoms yellow) of the X-ray structure (PDB-ID 4XXD ^13^. Hydrogen bonds are highlighted by dashed lines **B** Structural model of the Aβ_9-33_ A21C A30C monomer variant in the solanezumab binding pocket. The differences to the crystal structure are **1**. Elongation of the Aβ peptide up to residue 9-33. **2**. Mutation of residue A21 to C21 and formation of the intrapeptide disulfide bridge. **3**. Slight optimization of the crystal structure residues beginning by C21 to allow for this disulfide bridge. **C** Sequence alignment between the resolved Aβ wild-type residues in the X-ray structure and the monomer mutant, showing the position of the mutations and the disulfide bridge. Grey residues were present in the experiment but were not resolved.

### *Insights into* Aβ *peptide binding*

After confirming the theoretical predictions by the experiments, we performed a deeper theoretical analysis of the antibody-antigen binding. Two cognate antigens, Biotin-Aβ_1-40_ and the so-called monomer-mutant Aβ_9-33_ A21C A30C (Fig. 3A and 4), were modeled into the structure as detailed in the Methods section. A subsequent molecular dynamics simulation revealed that hydrophobic interactions, especially those of F19 and F20 to the antibody, are indispensable for the binding of the antigens. While additional contacts are frequently observed, they are often fluctuating in nature and even missing in some simulations, showcasing their supportive but not crucial nature.

Biotin-Aβ_1-40_, GST-Aβ_13-28_ (Fig. S1) and the monomer variant Aβ_9-33_ A21C A30C (Fig. 4) were modeled into the binding pocket using the experimentally resolved residues as a basis and extending the sequence. Slight steric hindrances were observed only for the substitution of A21C in the monomer variant (Fig. 4), which was resolved with a local optimization, as detailed in the Methods section.

To obtain dynamic insights into binding, Biotin-Aβ_1-40_ and the monomer variant bound to solanezumab were simulated in a water box with physiological salt concentration for 500 ns. The key binding interface is formed by the central Aβ residues 16-24 being tightly bound to the antibody is demonstrated by the Cα RMSD analysis (Fig. S2 and S3). These central residues deviate only little from the starting structure (∼1.5 Å) and are very rigid (fluctuation by ∼0.2 Å) during the simulation. The N- and C-terminal regions of both constructs, which extend outward from the binding pocket, display high fluctuations and show only sporadic contact with the antibody (Fig. S4+5).

The asymmetric unit cell comprises two copies of the complex, which superimpose with minimal differences (Cα RMSD of 0.21 Å). The primary distinction is observed for residues G25 and S26 of Aβ: these residues are resolved in chain C, where they engage in crystal packing contacts with symmetry-related molecules, but are disordered and not resolved in chain F in the absence of such stabilizing interactions. As G25 and S26 do not contribute to the antigen-antibody interface, they are omitted from Fig. 1 for clarity.

Detailed binding motives were identified via dynamic contact analysis (Fig. S4+5), which revealed the recognition of key contacts indispensable for the binding of both constructs. While polar contacts of the K16 and D23 sidechains, as well as the L17, F19 and A/C21 backbone to the antibody, were observed in parts of the simulation, their absence did not affect the hydrophobic core region, which consists of the sidechains of L17, V18, F19 and F20, whose embedding remained stable and unaffected by the surrounding fluctuations. Our approach to predict a dead mutant thus focused on disrupting the binding inside this core region.

### Application of the results for CSF measurements

Finally, we evaluated our system using cerebrospinal fluid (CSF), a sample type commonly used in clinical studies (Fig. 5). Consistent with the previous results, stable binding of the Aβ content of CSF to wild-type solanezumab was observed. Owing to the content of further proteins in CSF, the initial signal included both the contribution from bound Aβ and a bulk signal from unbound proteins circulating over the surface. Upon washing with buffer (indicated by the brackets in Fig. 5), the bulk signal disappeared, leaving only the signal from the bound Aβ in the case of the wild-type antibody. In contrast, no residual signal was observed for the mutant antibody, indicating an absence of binding. Thus there is also no unintended or unspecific binding from any compound of the CSF to the surface, demonstrating the good and robust performance of the iRS biosensor.

**Figure 5.**
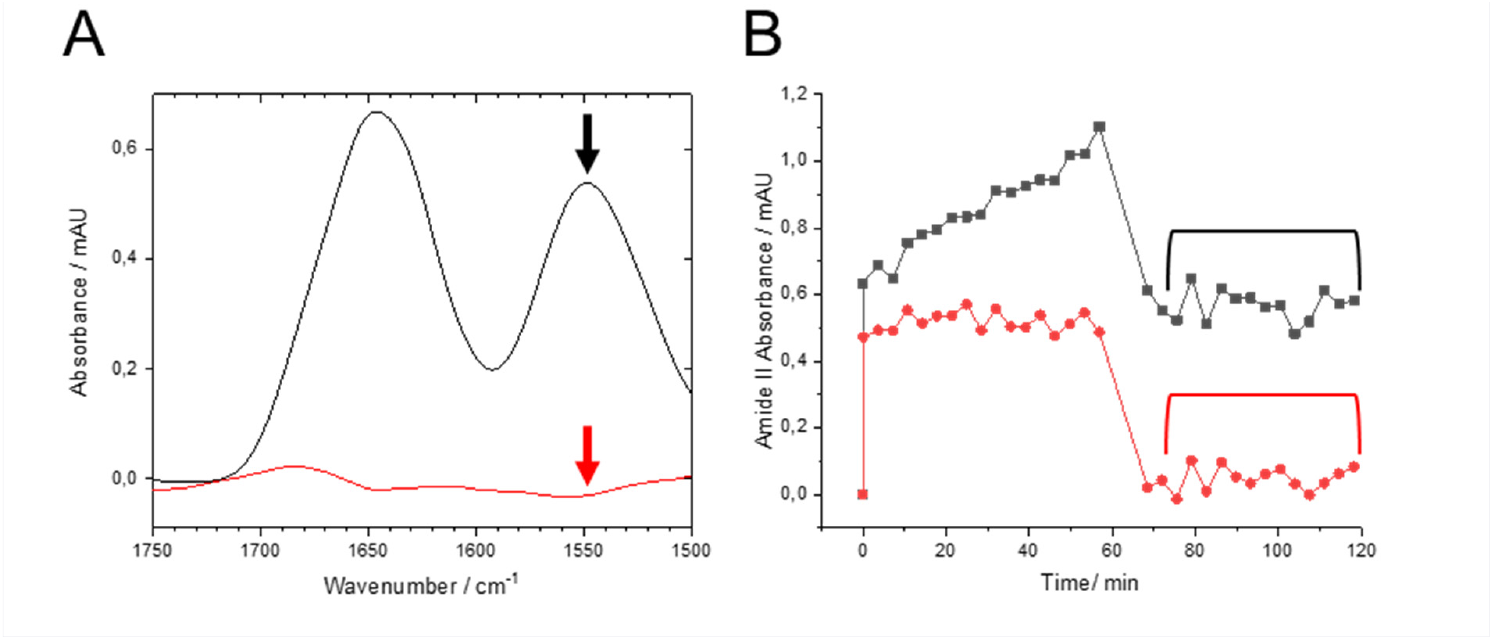
*In vivo* proof of the specificity of solanezumab binding. **A** ATR-FTIR measurements of cerebrospinal fluid (CSF) on a surface coated with the solanezumab antibody (black) or its G95A^HC^ dead mutant (red). It is evident that the dead mutant exhibits no binding, while the solanezumab wild-type antibody shows an amide II absorbance of 0.6 mAU. **B** Kinetics of amide II absorbance during CSF measurements for solanezumab wild-type (black) and the dead mutant (red) are presented. The amide II absorbance gradually increases over the 60-minute sample circulation in the wild-type, but remains constant in the dead mutant. Following the initiation of the wash step (indicated by the bracket), the amide II absorbance in the wild-type drops from 1 mAU to 0.6 mAU and remains stable throughout the wash step. Conversely, in the dead mutant, the entire amide II absorbance signal washes down from 0.5 mAU to 0. It was shown that the dead mutant does not bind Aß or any other compounds even from body fluids, demonstrating the specificity of the biosensor and the dead mutants ability to serve as a perfect negative control.

In line with our findings, a mutational analysis was performed by Ultsch et al. ^*26*^ for crenezumab, a highly similar antibody that targets the same Aβ epitope. In this study 14 single substitutions to alanine have been tested on their impact on binding affinity. The substitution positions were chosen based on their proximity to the Aβ, despite some of their side chains protruding away from the binding pocket. However, among the three variants that abolished Aβ binding as shown by SPR kinetic measurements is the G95A^HC^ substitution in crenezumab. Due to the structural similarity between crenezumab and solanezumab we anticipate the same structural rational for the “dead mutant” of crenezumab as for solanezumab.

## Conclusion

We demonstrate the power of combining computational protein modeling with experimental infrared-spectroscopic techniques to gain detailed insights into antibody-antigen interactions at the atomic level. By introducing a targeted mutation into the paratope of solanezumab, we were able to create a “dead mutant” that effectively abolished antigen binding without disrupting the overall structure of the antibody. We experimentally validated the dead mutant using pure antigens and for the complex body fluid CSF. The validation of the key interaction site allows further theoretical characterization of the binding. Experimentally, the “dead mutant” is a versatile tool with implications in diagnostic and possibly therapeutic applications. Our general workflow is applicable to any other antibody with its cognate antigen. In the absence of an experimental structural model, predicted structural models e.g. exploiting AI approaches, serve as basis for structural analysis and will then be validated by our strategy.

## Methods

### MD simulations

#### Protein Modeling

The crystal structure of solanezumab (PDB ID: 4XXD ^13^) served as the structural basis. It contains an asymmetrical unit cell with two copies of the antibody bound to Aβ. The experiment contained the entire Aβ_1-40_, of which only residues 16-26 in copy 1 and 16-24 in copy 2 were resolved, indicating an unstructured nature and no specific contacts of the remaining Aβ. Furthermore, residues 25 and 26 in copy 1 make no contact with the antibody itself and are stabilized only by crystal contacts with the surrounding unit cells. As such, residues 16-24 were taken as the basis for all the following modeling approaches performed with the MAXIMOBY/MOBY protein modeling software package ^27^.

Two structures were prepared for the molecular dynamics simulation: solanezumab bound to (a) wild-type Aβ_1-40_ and (b) Aβ_9-33_ A21C A30C, a double cysteine mutant forming a disulfide bridge, which serves as a non-aggregating monomeric mimic of Aβ in experiments. Due to missing structural information, the remaining residues 1-15 and 25-40 for construct (a) were modeled in an all-trans conformation and subsequently energy optimized utilizing the united atom AMBER forcefield ^28^.

For construct (b), A21 was mutated to cysteine within MOBY. Due to a steric clash of the newly introduced cysteine side chain with Y27^LC^ of the antibody, the backbone and side chain of C21 were locally optimized. To fully resolve the clash, adjacent residues 22-24 were also refined, resulting in a slightly altered local conformation while preserving all key contacts (Fig. 4).

The Aβ N-terminus was again modeled in an all-trans conformation. The C-terminal region was similarly modeled in an all-trans backbone conformation up to residue 30, where the second mutation (A30C) was introduced. A disulfide bridge between C21 and C30 was then formed, and residues 25–30 were adjusted to accommodate the new covalent linkage. Finally, residues 31–33 were appended to complete the model. A comparison of the resulting structures is shown in Fig. 4.

Two additional structures were modeled, following the same principles of minimal intervention in the experimental structures: solanezumab bound to (c) GST-Aβ_13-28_ and (d) Biotin-Aβ_1-40_.

#### Molecular Dynamics Simulations

Molecular dynamics (MD) simulations are prepared and analyzed with the MAXIMOBY/MOBY protein modeling software package ^27^ and performed with the GROMACS molecular dynamics program suite version 2022 ^29^. To prepare the protein structure, the first step is an atom wise analysis of the potential energy. For this purpose, the united atom AMBER force field ^28^ is chosen. The united atom version of AMBER is employed, as many experimental structures do not contain hydrogen atoms. Energetically unfavorable side chain and backbone conformations are optimized through energy minimization of as few atoms as possible to resolve the issue. This was only needed for the substitution of A21C in the Aβ_9-33_ A21C A30C, as described above.

Next, the protonation states of amino acids are determined on the basis of the pKa calculation of Nielsen & Vriend ^30^. Every protonable amino acid side chain or protein terminus is assigned a standard pKa value, which is then further influenced by its local surroundings, as calculated via the QEq method^31^. The targeted pH value for the simulation system was set to seven. Every calculated pKa value is compared to this value to determine the final protonation state of the respective functionality. To finalize protonation, the functionalities whose pKa is closest to their neutral state are systematically decharged to obtain a total charge of zero for the system. For residues with more than one protonable atom, such as histidine, the hydrogen is placed in a manner that allows the best hydrogen bond interaction with the surrounding.

The orientation of the nitrogen and oxygen in the ASN and GLN side chains is not determinable through most experimental structure resolving methods. Because of this, both possible orientations are analyzed on the basis of their ability to form hydrogen bonds with the surrounding residues. The best orientation is subsequently chosen.

Using the solvation procedure implemented in MAXIMOBY, which is based on the Vedani algorithm ^32^, water molecules of the first and second protein solvation shells are placed.

A cubic simulation box with a minimum distance of 1.4 nm to the outer atoms of the second solvation shell is placed around the protein. This choice of dimensions prevents the long-range interaction of water molecules of the second solvation shell from one side of the protein to the other due to the periodic boundary conditions using a cutoff of 1.1 nm. The remaining simulation box is filled with TIP4P water ^33^ using the solvation algorithm implemented in GROMACS. A physiological NaCl concentration of 0.154 mol/l was also added with GROMACS. The algorithm used replaces water molecules randomly with either NA^+^ or Cl^-^. We have fine-tuned this step by not replacing any water molecules of the two solvation shells, which prevents ions from being inserted inside or close to the protein in the starting structure and thus possibly destabilizing it. The solvated system is again checked for energetic clashes and locally optimized if needed. The solvation algorithm of GROMACS does not place water molecules on the basis of hydrogen bonds but rather fills all available spaces. Because of this, an optimization of all water hydrogens is performed here to ensure a smooth transition between the solvation shells and the bulk water added by GROMACS, as well as between ions and water molecules.

All the following steps are performed with the Optimized Potential for Liquid Simulations (OPLS)-All-Atom forcefield ^34^, a continuation of the AMBER forcefield with fine-tuned nonbonded parameters to reproduce correct experimental values in dense phase (liquid) simulations.

First, the system is heated to 298 K over 1 ns. The temperature is increased linearly from 0 to 100 K during the first 0.1 ns. The remaining 0.9 ns are used to linearly heat the system to the desired 298 K. The heating is performed with a timestep of 1 fs. A velocity-rescale Berendsen thermostat is used, with the temperature coupling updated every 100 fs. We define two temperature coupling groups: 1. The protein and the two solvation shells, 2. The bulk water and ions. The bond lengths and angles of Tip4p water are constrained by the SETTLE algorithm ^35^. This leads to a lower energy capacity and thus would lead to a faster heating process, compared to the protein, which is why the bulk water molecules and ions are coupled separately. The following equilibration and production runs are performed with a timestep of 2 fs. To use this timestep, all h-bond lengths are constrained by the LINCS algorithm ^36^.

Pair interactions use the grid-based Verlet scheme ^37^ with a Coulomb and vdW cutoff of 1.1 nm. The vdW interactions use a switching function, which scales the interaction energies linearly to zero starting at 1.05 nm and ending at the cutoff. Coulomb interactions are calculated via a fourth-order PME scheme with a Fourier spacing of 0.12 nm. Finally, the center of mass translational movement is removed every ps for the whole system.

For equilibration, a 1 ns nVT run with constant particle number (n), volume (V) and temperature (T) without pressure coupling (p) is performed. New velocities are generated at the start of the nVT run. Thus, it is the starting point when multiple runs of the same system are performed. The temperature is controlled with the velocity-rescaling thermostat to ensure the proper canonical ensemble ^38^. The temperature is coupled at every step with a time constant (tau-t) of 0.1 ps.

This run is followed by a 10 ns npT run with a constant particle number (n), pressure (p), and temperature (T) but a flexible volume (V). The temperature and pressure are controlled every V-rescale thermostat and a Berendsen barostat with a time constant (tau) of 0.1 ps ^39^. The target pressure is set to 1 bar with an isotropic compressibility of 4.5*10^−5^ bar^-1^.

The final npT production run uses the Nose-Hoover thermostat ^40^ and Parinello-Rahman barostat ^41^ with a tau-t of 0.5 ps and a tau-p of 2.5 ps.

#### Simulation Evaluation

To assess the stability of the simulation, the root mean square deviation (RMSD) of the Cα atoms was calculated for each snapshot of the trajectory relative to the starting structure. Changes in the secondary structure were monitored using the DSSP (Define Secondary Structure of Proteins) algorithm. Inter-protein interactions were analyzed using PyContact^14^ and the contact matrix algorithm implemented in MAXIMOBY. Hydrogen bonds were defined based on distance and angle criteria between donor and acceptor atoms and further classified into backbone-and side chain-derived contributions. Non-specific van der Waals (vdW) contacts were identified by carbon–carbon distances, while specific vdW interactions involving π-systems were additionally assessed based on the relative spatial orientation of the interacting partners.

#### ATR-FTIR spectroscopy

We employed an enhanced functionalization method as outlined in patents WO2024003213A1 and WO2024003214A3 ^16,17,42^, which will be elaborated upon in a separate publication. These enhancements resulted in significantly improved stability and inertness of the ATR surface compared with previous reports. The initial linker molecule incorporates a triethoxysilane functional group, enabling covalent attachment to the silicon ATR surface. Subsequent modification steps utilize either strain-promoted azide-alkyne cycloaddition (SPAAC) chemistry, specifically the reaction of azadibenzocyclooctyne (DBCO) with azide groups ^43,44^, or the coupling of N-hydroxysuccinimide (NHS) esters with primary amines ^45^. All functionalization steps were conducted within the flow-through system of the spectrometer, permitting real-time monitoring of each reaction. The resulting surface architecture comprises a covalently immobilized blocking layer, to which the catcher molecule is also covalently attached. For surface immobilization, the E7cys454 was site-directedly coupled with the EZ-Link™ maleimide-PEG4-DBCO-linker(*Thermo Fisher Scientific, Waltham, USA*) according to the manufacturer’s instructions. After surface preparation, the crystal surface was functionalized with E7 (50 μg in 50 mM MES + 0.05% Tween buffer, pH 5.5). Subsequently, solanezumab (wild-type or dead mutant)−Im7 (40 µg in MES buffer, pH 5.5) was immobilized on the surface. In the next step, the samples, consisting of two Aß antigens (1 µg 200 ng/µl GST-Aß_13-28_ and 0.5 µg 200 ng/µl Aβ_9-33_ A21C A30C) or 300 µl CSF, were circulated over the surfaces for one hour in PBS buffer at pH 7.4. This was followed by a 30-minute wash step with PBS pH 7.4 buffer. All surface processes were monitored by recording the corresponding infrared spectra at each step, enabling precise control over each reaction stage. The silicon-IRE was integrated into a Fourier transform infrared (FTIR) Vertex 80 V spectrometer (Bruker Optics, Ettlingen, Germany) in conjunction with a liquid nitrogen-cooled mercury cadmium telluride (MCT) detector and a mid-infrared (MIR) source. The device setup and spectrometer parameters for spectral acquisition have been previously described in detail ^15^. The ATR unit (Specac Ltd., Slough, UK) was installed in the sample compartment of the FTIR instrument and aligned at an incidence angle of 45°. Measurements were conducted under flushing with dry air to eliminate the sharp atmospheric water vapor absorbance bands. The CSF samples used are surplus material from MVZ Labor Leipzig, Leipzig, Germany. They are anonymized and cannot be traced back to the donor.

#### Expression of E7cys454

The gene encoding Colicin E7 (UniProt: Q47112) was cloned into a pET24a plasmid. A T454C substitution (threonine to cysteine) was introduced into the amino acid sequence to enable site-specific conjugation. Further, histidine 545 was mutated to alanine in order to abolish DNase activity^46^. Competent *E. coli* BL21-AI™ (One Shot™) cells were transformed with 100 ng of plasmid DNA and plated on LB agar supplemented with kanamycin (50 µg/mL). Multiple colonies were used to inoculate 10 mL LB pre-cultures containing kanamycin (50 µg/mL), which were incubated overnight at 37°C with shaking (220 rpm). These cultures were then used to inoculate 5 L of Terrific Broth (TB) medium to an initial OD600 of 0.05. Cells were grown at 37°C with agitation (80 rpm) until reaching mid-log phase (OD600 0.6-0.8). Protein expression was induced by adding 0.02% (w/v) L-arabinose and 1 mM isopropyl β-D-1-thiogalactopyranoside (IPTG), followed by incubation at 25°C for 16 hours. After overexpression was complete, the cells were spun down (5000 × g, 15 minutes, 4°C) and resuspended in 10 ml/l PBS (pH 7.4). The cells were then disrupted via five passages through a microfluidizer (1000 bar pressure). The resulting material was centrifuged (45000 × g, one hour, 4°C) and separated into two components: the soluble fraction in the supernatant (cytoplasmic fraction) and the insoluble fraction in the sediment. The supernatant was then filtered (pore size of 0.2 μm) and purified via affinity chromatography column. Purification was carried out using the His-Tag (HisTrap™), which is attached to the heavy chain, followed by purification using size exclusion chromatography.

#### Expression of the solanezumab-Im7 wild-type and the solanezumab-Im7 dead mutant

The solanezumab-Im7 fusion proteins used in this study were modified on the basis of the publication by Hosse et al. and cloned as a bicistronic vector construct in the PET24a plasmid ^18^. For this purpose, the Im7 gene (UniProt: Q03708) was fused to the 3’ end of the solanezumab light chain via an intermediate flexible peptide linker, (G4S)2-GGRAS (Fig. S6). The solanezumab wild-type Im7 fusion protein under investigation, as well as the proposed inactive mutant G95A, were produced under identical conditions through protein overexpression in *E. coli*. For this purpose, the respective plasmid DNA was transformed into *E. coli BL21-AI™ (One Shot™*) cells, which were subsequently grown in LB precultures with kanamycin (50 µg/ml) overnight at 37°C with shaking. For expression, 5 L of TB medium was inoculated at an OD600 of 0.05 and induced at an OD600 of 0.6-0.8 with 0.02% arabinose and 1 mM IPTG. From the time of induction, the temperature was reduced to 18°C, and the cultures were incubated for 40 hours with constant shaking. After overexpression was complete, the cells were spun down (5000 × g, 15 minutes, 4°C) and resuspended in 10 ml/l PBS (pH 7.4). The cells were then disrupted via five passages through a microfluidizer (1000 bar pressure). The resulting material was centrifuged (45000 × g, one hour, 4°C) and separated into two components: the soluble fraction in the supernatant (cytoplasmic fraction) and the insoluble fraction in the sediment. The supernatant was then filtered (pore size of 0.2 μm) and purified via two affinity chromatography columns. First, purification via the His-tag (HisTrap™) attached to the heavy chain was performed, and then, purification via the Strep tag (StrepTrap™ HP) of the light chain was performed. The combined Fab fusion protein contains both tags and is ultimately purified by size exclusion chromatography.

#### Expression of GST-Aβ_13-28_

To characterize our Im7-Fab fusion proteins, a glutathione S-transferase (GST)-Aß_13-28_ fusion protein was produced in *E*.*coli*. For this purpose, the GST from *S. japonicu*m was fused n-terminally to the Aß_13-28_ peptide to form a conjugate. This gene was produced in *E. coli* by overexpression. For this purpose, the respective plasmid DNA was transformed into *E. coli BL21-AI™ (One Shot™*) cells, which were subsequently grown in LB precultures with kanamycin (50 µg/ml) overnight at 37°C with shaking. For expression, 5 L of TB medium was inoculated at an OD600 of 0.05 at 37°C and induced at an OD600 of 0.6--0.8 with 0.02% arabinose and 1 mM IPTG. From the time of induction, the temperature was reduced to 30°C, and the cultures were incubated for 16 hours with constant shaking. After overexpression was complete, the cells were spun down (5000 × g, 15 minutes, 4°C) and resuspended in 10 ml/l PBS (pH 7.4). The cells were then disrupted via five passages through a microfluidizer (pressure 1000 bar). The resulting material was centrifuged (45000 × g, one hour, 4°C) and separated into two components: the soluble fraction in the supernatant (cytoplasmic fraction) and the insoluble fraction in the sediment. The supernatant was then filtered (pore size of 0.2 μm) and purified via affinity chromatography. For purification, we used GST affinity chromatography (GST HiTrap®), in which the GST fusion protein is bound to the column via glutathione Sepharose. This was followed by size exclusion chromatography for final purification.

#### ELISA Biotin-Aß_40_

The ELISA was based on previously described methods for the analysis of Fab fragments using ELISA ^24^. For the ELISA, biotin-conjugated Aß_1-40_ (Biotin-Aß_1-40_) was bound to streptavidin-coated plates (*Thermo Fisher Scientific, Waltham, USA*). For this purpose, 1 µg/ml biotin-Aß_1-40_ was incubated in assay buffer (PBS with 1% bovine serum albumin) at 50 µl/well for one hour. The plates were then washed with wash buffer (PBS with 0.05% Tween-20) and blocked for one hour with blocking buffer (PBS with 1% BSA). The Fab fusion fragments were incubated on the plate in a serial dilution series for two hours. The samples were subsequently washed three times with wash buffer and incubated with an HRP-conjugated polyclonal anti-human secondary antibody. Detection was achieved through the enzymatic reaction of horseradish peroxidase with the substrate O-phenylenediamine dihydrochloride (OPD) (Thermo Fisher Scientific, Waltham, USA). The absorbance was measured at 492 nm via a CLARIOstar® Plus plate reader (BMG Labtech, Ortenberg, Germany).

## Acknowledgment

The presented research was funded by the Center for Protein Diagnostics (PRODI), Ministry of Culture and Science of North-Rhine Westphalia, Germany. We thank Janine Trautmann and Dr. Thorsten Klemm for providing the CSF samples. We thank PD Dr. Udo Höweler for fruitful discussions regarding the antigen antibody interaction data.

