## Supplemental Figures S1-S6 for "Replacement of a single residue in an antibody completely abolishes cognate antigen binding, as predicted by theoretical methods"

**Figure S1. Structural models of monomer mimicking Aβ constructs and their binding to solanezumab. A** Predicted structural model of the fusion complex of GST (gray cartoon representation) with Aβ 13-28 (sticks representation of carbon and nitrogen backbone atoms) bound to the antibody (surface representation, heavy chain: orange, light chain: green). The binding interface was modeled based on the X-ray structure with PDB-ID 4XXD (Crespi et al. 2015) The experimentally resolved Aβ residues 16-26 (carbon atoms teal, nitrogen atoms blue) are expanded N-terminally by the Aβ residues 13-15 and C-terminally by residues 27-28 (cyan carbon atoms), the Strep-Tag at the C-terminus (gray carbon atoms), and the linker to the GST (gray carbon atoms). **B** Structural model of Aβ 1-40-Biotin bound to solanezumab modeled analogous to the GST-Aβ fusion complex described in A. Here the Aβ-peptide is expanded by residues 1-15 and 27-40 as well as by the N-terminally added biotin. The structural models do not indicate any significant differences in the expected antigen binding affinities between the different constructs. This expectation was confirmed by our experimental measurements shown in Fig. 3.

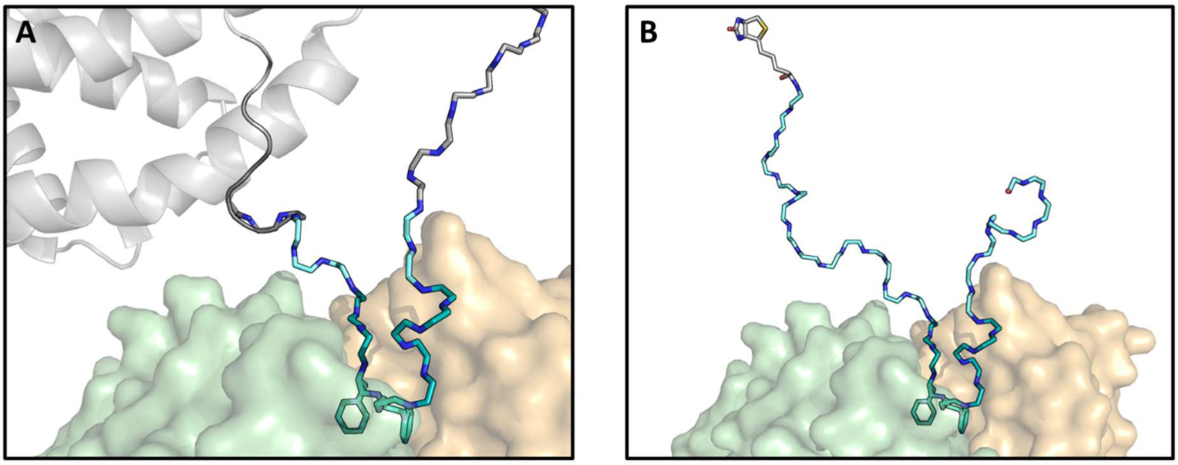


MSPILGYWKIKGLVQPTRLLLEYLEEKYEEHLYERDEGDK
WRNKKFELGLEFPNLPYYIDGDVKLTQSMAIIRYIADKHN
MLGGCPKERAEISMLEGAVLDIRYGVSRIAYSKDFETLKV
DFLSKLPEMLKMFEDRLCHKTYLNGDHVTHPDFMLYDALD
VVLYMDPMCLDAFPKLVCFKKRIEAIPQIDKYLKSSKYIA
WPLQGWQATFGGGDHPPKLVPRGSHHQKLVFFAEDVGSNK
WSHPQFEK

Biotin-DAEFRHDSGYEVHHQKLVFFAEDVGSNKGAIIG
LMVGGV

**Figure S2. C$\alpha$ RMSD for the simulations of A$\beta$1-40 bound to solanezumab.** Regarding the RMSD, all structures are stable after about 150-300 ns. The RMSD for the A$\beta$ was subdivided into two parts: Residues 16-26, which are resolved in the crystal structure 4xxd were calculated separately from the modeled residues 1-15 + 27-40. As the latter ones were not resolved, they were modeled in an all-trans conformation and consequently show a high deviation from the starting structure. However, the RMSD also shows higher fluctuations, indicating that these parts are flexible and thus make no specific contacts to the antibody. This was confirmed in the contacts analysis (Fig. S4). The central part of A$\beta$ is tightly positioned in the binding pocket, reflected in its stable RMSD of 1-2 Å.

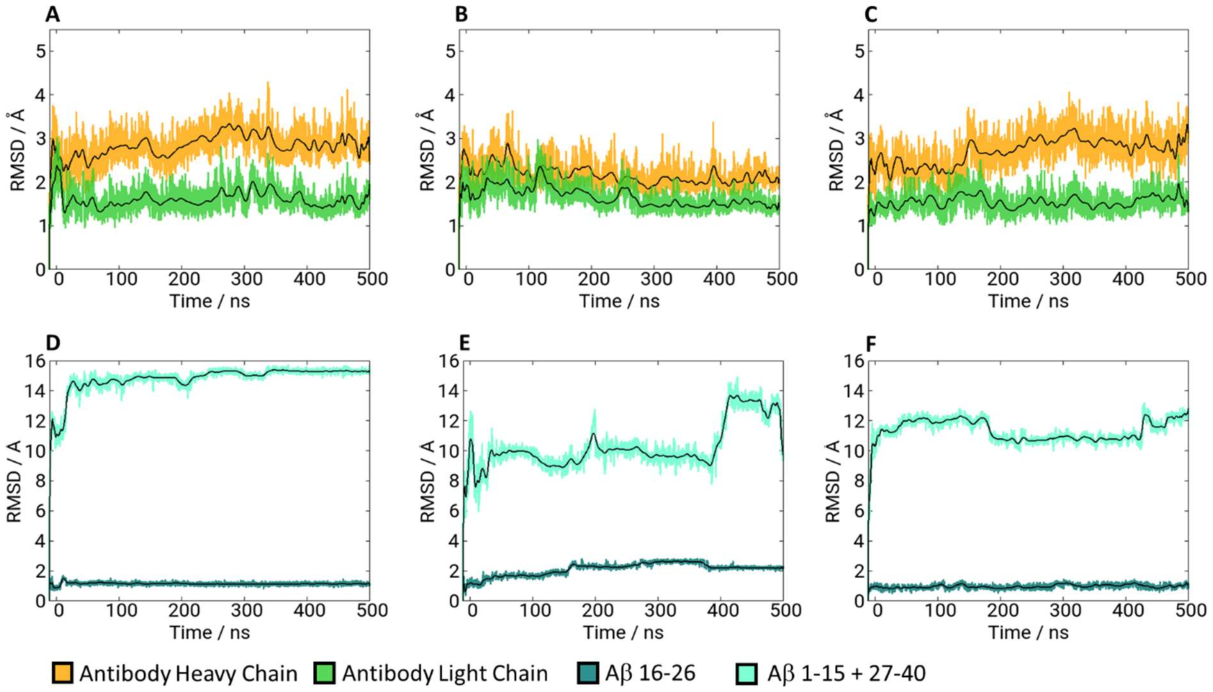

**Figure S3. Cα RMSD of the simulations of Aβ9-33 A21C A30C bound to solanezumab.** Regarding the RMSD, all structures are stable after about 200 ns. The RMSD for the Aβ was subdivided into two parts: Residues 16-26, which were closely modeled to the resolved Aβ residues of the crystal structure 4xxd were calculated separately from the residues 9-15 + 27-33. As the latter ones had no equivalent in the experimental structure, they were modeled in an all-trans conformation and consequently show a high deviation from the starting structure. However, the RMSD also shows higher fluctuations, indicating that these parts are flexible and thus make no specific contacts to the antibody. This was confirmed in the contacts analysis (Fig. S5). The central part of the Aβ mutant is tightly positioned in the binding pocket, reflected in its stable RMSD of 1-2 Å.

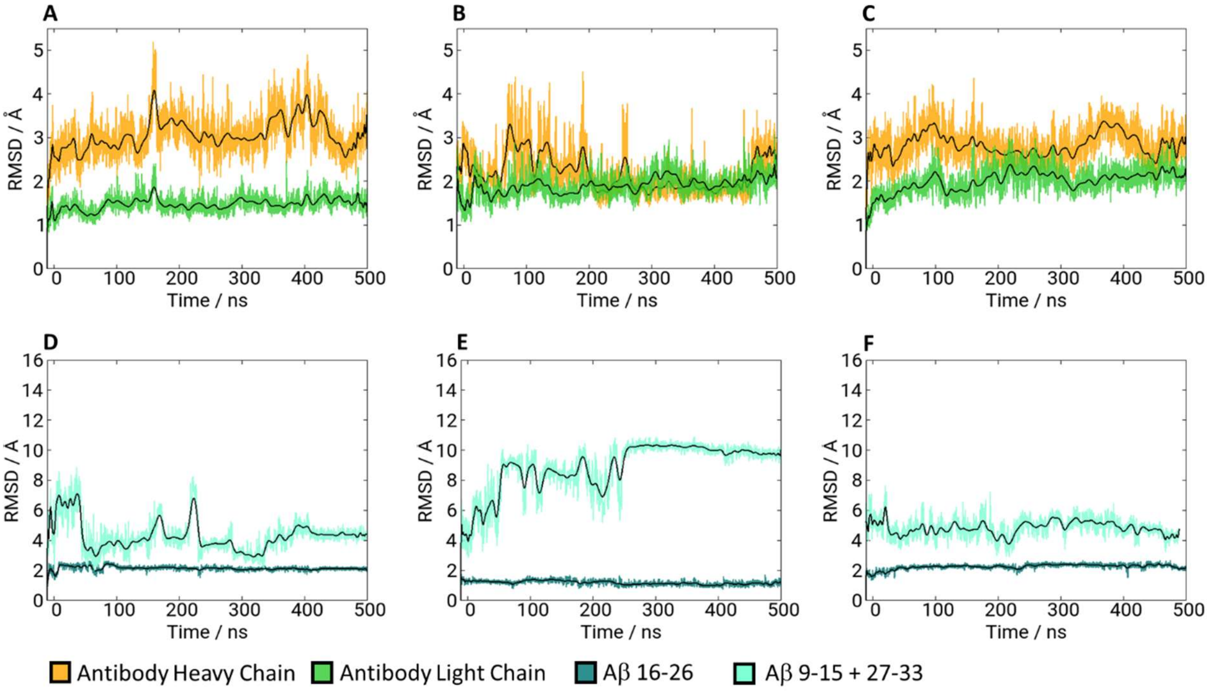

**Figure S4. Contact plots for the simulations of Aβ1-40 bound to solanezumab.** Shown are all contacts between the respective Aβ peptide and the antibodies (Aβ = chain A, antibody heavy chain = chain H, antibody light chain = chain L), which are present either present in the crystal/starting structure or for at least 50 % of the second half of the simulation, therefore in the fully equilibrated structure. The dynamic analysis highlights the importance of hydrophobic contacts (blue color scheme) for the binding of Aβ by solanezumab. Polar contacts (yellow, orange and red) are not essential for the binding and are frequently fluctuating. The first and second column indicate the presence of the contact in the starting structure and after the equilibration. Starting at 0 ns, the presence during the production run is shown with a contacts analysis every 0.1 ns.

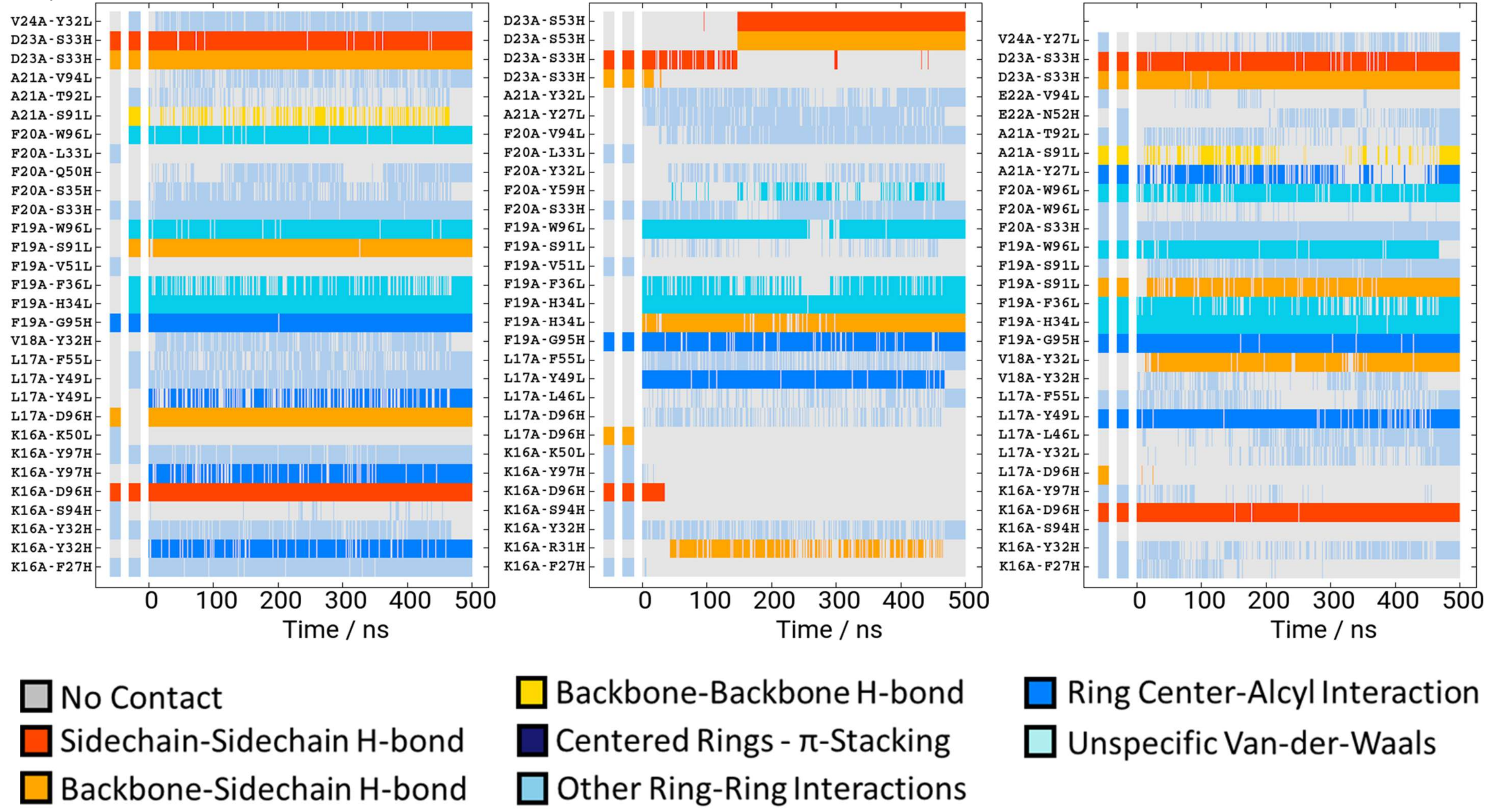

**Figure S5. Contact plots for the simulations of Aβ9-33 A21C A30C bound to solanezumab.** Shown are all contacts between the respective Aβ peptide and the antibodies (Aβ = chain A, antibody heavy chain = chain H, antibody light chain = chain L), which are present either present in the crystal/starting structure or for at least 50 % of the second half of the simulation, therefore in the fully equilibrated structure. The dynamic analysis highlights the importance of hydrophobic contacts (blue color scheme) for the binding of Aβ by solanezumab. Polar contacts (yellow, orange and red) are not essential for the binding and are frequently fluctuating. The first and second column indicate the presence of the contact in the starting structure and after the equilibration. Starting at 0 ns, the presence during the production run is shown with a contacts analysis every 0.1 ns.

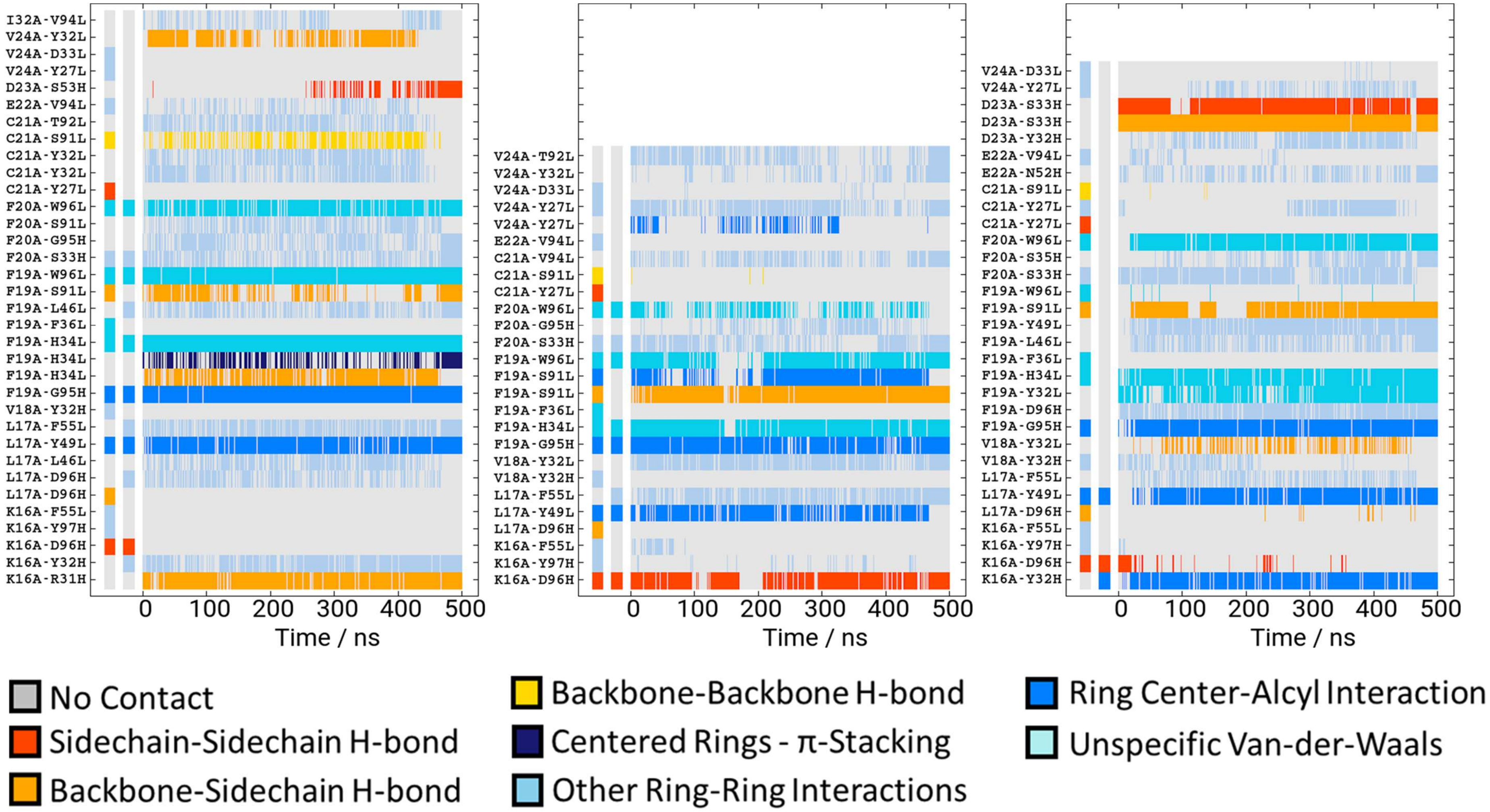

**Figure S6: Extract of the bicistronic vector construct for the recombinant expression of solanezumab-Fab-Im7 in E. coli**. The red areas code for the Fab-fragment of solanezumab. Attached to the light chain gene is the genetic information for Im7 and a connecting flexible linker is cloned to the light chain gene.

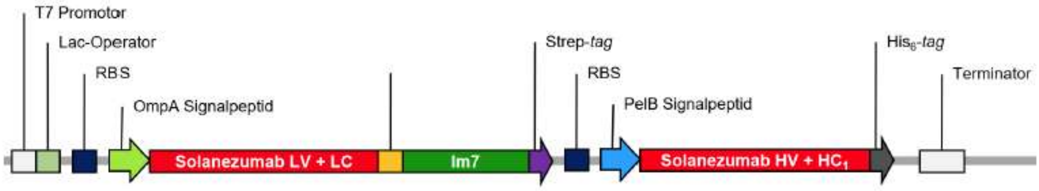
